# Consolidation of behavioral synchronization with the developmental clock is impaired by the degeneration of a mechanosensory circuit in *C. elegans*

**DOI:** 10.1101/2024.10.28.620731

**Authors:** Lior Laufer, Sharon Inberg, Eshkar Nir, Adi Sufrin, Shay Stern

## Abstract

Animals generate predictable patterns of behavior that are robustly synchronized with the developmental clock. However, the dynamics of the long-term establishment of synchronized behaviors across the entire developmental trajectory, and the neuronal circuits that control this developmental synchronization, remain underexplored. Here, we show that in *C. elegans* individuals, developmental synchronization of behavior continuously consolidates throughout development time, and that this consolidation is controlled by multiple neuronal pathways. In particular, we found that the degeneration of a specific mechanosensory circuit early in development strongly impairs the developmental synchronization of behavior. Furthermore, early neuronal activity patterns in downstream motor neurons are altered following the mechanosensory circuit degeneration. Interestingly, we show that either pharmacological or genetic neuroprotection of the mechanosensory circuit from degeneration restores normal modes of developmentally synchronized behaviors. These results imply the control of time-locked behaviors across the developmental trajectory by a localized sensory circuit within the nervous system.

## Introduction

Individuals exhibit patterns of behavior over long timescales that are temporally synchronized. For example, activity and sleep cycles are regulated by a circadian rhythm across species (Konopka and Benzer, 1971; Panda et al., 2002). In addition, the lunar cycle affects the foraging of fruit bats and kangaroo rats (Daly et al., 1992; Morrison, 1978), and circannual rhythms are associated with bird migration (Gwinner, 1986), as well as hibernation cycles in chipmunks (Kondo et al., 2006). In particular, during development, animals display predictable temporal changes in behavior that depend on the specific developmental window and are synchronized with the developmental clock of each individual. For instance, *Drosophila melanogaster* shows a ‘foraging’ state during the third larval stage and a ‘wandering’ state during the pre-pupation stage (Sokolowski et al., 1984). In addition, behavioral patterns in honeybees are modified over their lifetime (Vance et al., 2009), and the startle response in fish is shaped throughout development (Kimmel et al., 1974). Moreover, fear extinction learning in both humans and mice is altered at specific developmental windows (Pattwell et al., 2012).

*C. elegans* is an efficient model system for studying the temporal synchronization of behavior across developmental timescales due to its short development time of 2.5 days, genetically homogeneous population, and the well-defined structure of its nervous system (White et al., 1986). It was previously demonstrated that isolated *C. elegans* individuals show stereotyped long-term patterns of behavior that are highly dynamic throughout development (Stern et al., 2017; Harel et al., 2024). However, it is still unclear how behavioral synchronization with the developmental clock is continuously established across and within all developmental stages, and which circuits within the nervous system are required for the robust time-synchronization of behavior across the entire developmental trajectory.

The function of the nervous system can be shaped by the modification of neuronal states, as well as by structural changes within the nervous system during development or adulthood. In *C. elegans*, both naturally occurring programmed cell death via apoptosis (Ellis and Horvitz, 1986) or neurodegeneration via cellular necrosis (Chalfie and Wolinsky, 1990; Driscoll and Chalfie, 1991; Treinin and Chalfie, 1995) were shown to modify the nervous system structure during its formation. Here, by monitoring the long-term locomotory behaviors of multiple isolated *C. elegans* individuals across their full development time and quantifying their developmental synchronization of behavior, we found a gradual consolidation of temporal behavioral synchronization across developmental stages. We further revealed that multiple neuronal pathways may either increase or decrease the degree of developmental synchronization of behavior.

Interestingly, we found that neurodegeneration of a specific mechanosensory circuit strongly impairs the robustness of temporal synchronization of behavioral patterns across development. At the neuronal activity level, the degeneration of the localized mechanosensory circuit led to alterations in the activity dynamics of downstream motor neurons early in development. We further showed that neuroprotection of the mechanosensory circuit from degeneration can reestablish normal modes of long-term synchronization of behavior. Overall, these results uncover the establishment and circuit control of robust temporal synchronization of behavioral patterns with the developmental clock.

## Results

### *C. elegans* shows consolidation of behavioral synchronization with the developmental clock across stages

To study the temporal organization of behavior across development at the individual and population level, we utilized long-term behavioral monitoring of multiple individuals across their complete developmental trajectory (Stern et al., 2017; Harel et al., 2024). In particular, in this imaging system, the behavior of single isolated individuals is continuously tracked from egg hatching to 16 hours of adulthood (∼2.5 days) in custom-made multi-well plates, at high spatiotemporal resolution (3 fps, ∼10 µm) and under a tightly controlled environment (Fig. 1A). Wild-type individuals show variation in their development time (±5%) (Fig. S1A). To align behavioral trajectories of different individuals we divided the developmental stage of each single individual into 75 time windows (Fig. S1B) (Stern et al., 2017; Ali Nasser et al., 2023). While growing in a food environment, *C. elegans* shifts between active and inactive locomotory states that last seconds to minutes, called roaming and dwelling (Ben Arous et al., 2009; Flavell et al., 2013; Fujiwara et al., 2002; Stern et al., 2017). It was previously shown that during development, individuals exhibit long-term patterns of roaming and dwelling behavior that are highly dynamic across and within stages (Fig. 1A,C; Fig. S1C,D) (Stern et al., 2017). These long-term behavioral patterns are manifested by specific developmental periods of high roaming activity, such as during the first half of the L2, L3, and L4 larval stages, as well as peaks of roaming that are more limited in time, such as during the end of the L4 stage. These stereotyped temporal patterns of behavior shown among isolated individuals suggest that specific behavioral outputs in *C. elegans* are continuously synchronized with the developmental clock of each single individual. To systematically study this developmental synchronization, and how it is continuously established as development progresses, we quantified the behavioral time-correlation among wild-type individuals at distinct developmental stages. High behavioral time-correlations would indicate that different individuals within the population tend to show similar temporal patterns of roaming activity across developmental windows, indicating the robust continuous synchronization of long-term behavior with the time of development (Fig. 1B; Fig. S1E). In contrast, low time-correlations among individuals would indicate that the behavioral patterns of different individuals is variable in time (up to randomness), suggesting a low level of robustness in forming predictable patterns of behavior during specific developmental windows (Fig. 1B; Fig. S1E). By first quantifying the temporal synchronization among wild-type individuals across their complete developmental trajectory, we found that overall, individuals showed higher behavioral synchronization over time, compared to a randomly shuffled dataset (Fig. S1F). Next, to further study whether the levels of developmental synchronization of behavior change across distinct developmental stages, we separately analyzed behavioral time-correlations among individuals within each of the five developmental stages (L1 to L4 and adulthood). Interestingly, we found that the robustness of developmental time-synchronization increased across larval stages (Fig. 1C,D). In particular, during the L1 stage, individuals showed patterns of roaming activity that are disorganized in time, reflected by the close-to-random correlations during this stage. Following the transition to the L2 stage, individuals showed significantly higher temporal synchronization, which gradually increased during the L3 and L4 stages (Fig. 1C,D). After the transition to adulthood, animals showed a slight decrease in temporal correlations, that were still significantly higher than in the early L1 stage. This strengthening of behavioral time-correlations across development time uncovers a ‘consolidation’ of developmental synchronization of behavior (Fig. 1D). These results reveal the continuous establishment of time-locked behaviors across the developmental trajectory of *C. elegans*.

**Figure 1.**
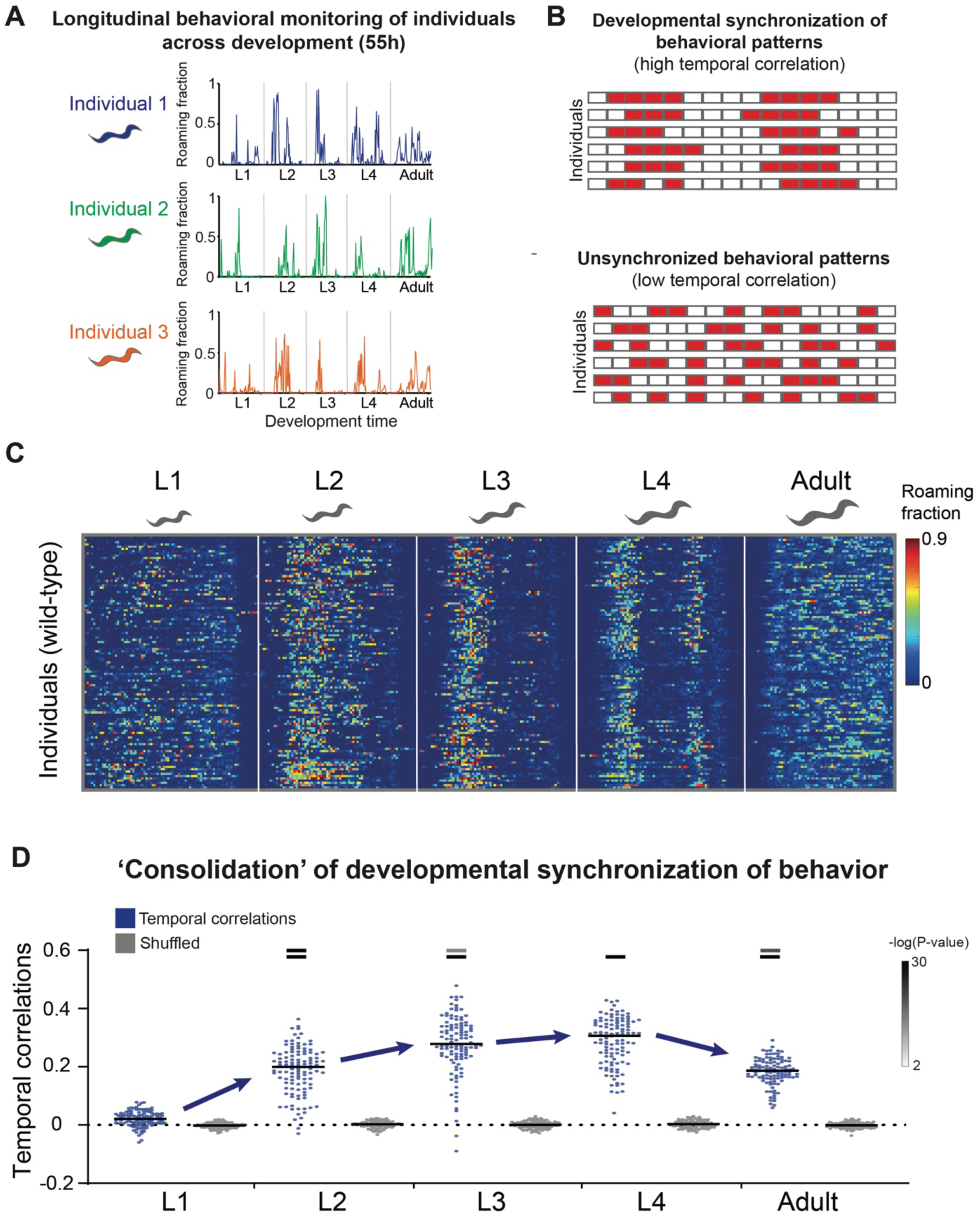
Long-term tracking of *C. elegans* individuals reveals the consolidation of developmental synchronization of behavior. **(A)** Shown are representative developmental trajectories of roaming behavior of three individual animals across all four larval stages and adulthood. Behavioral monitoring was performed by a multi-camera imaging system that allows longitudinal behavioral tracking of multiple *C. elegans* individuals throughout a complete development time, at high spatiotemporal resolution and under tightly controlled environmental conditions. Age-normalization equally divides each developmental stage into 75 time-bins. **(B)** A scheme representing temporally synchronized (top) and unsynchronized (bottom) behavioral patterns of roaming activity. Each row represents a single individual. Red indicates bouts of active roaming behavior over time. **(C)** Roaming and dwelling behavior of wild-type N2 animals (n=113). Each row indicates the age-normalized behavior of one individual across all developmental stages. Different developmental stages are separated by white lines indicating the middle of the lethargus state. Color bar represents the fraction of time spent roaming in each of the time-bins. **(D)** Temporal correlations of roaming behavior among individuals within the wild-type population (blue) compared to a randomly shuffled dataset (grey) across distinct developmental stages. Each dot represents the average temporal correlation between a single individual and all other individuals in the population. Black bars represent the median correlation. Color bars represent the significance (-log(P-value)) of the difference in temporal correlations between each of the L2-Adult stages to the L1 stage (bottom), and to the previous stage (top) (Wilcoxon rank-sum test, FDR corrected). Indicated are P-values<0.01.

### Identification of neuronal pathways affecting behavioral time-synchronization across development

To reveal underlying neuronal pathways that may affect the developmental synchronization of behavior, we performed a candidate screen to identify mutants in which the synchronization of behavior across developmental stages is altered. In particular, we systematically analyzed the developmental synchronization within 13 populations mutant for various neuronal pathways involved in neuromodulation, sensation, epigenetic regulation, and modifications of circuits within the nervous system (Fig. 2A; Fig. S3A). For each population, we quantified the temporal correlations in each of the developmental stages and compared them to the wild-type population. These analyses revealed multiple effects on the developmental synchronization of behavior that may be systematic across most of the developmental trajectory or exposed at specific stages. For instance, individuals mutant for the epigenetic regulators *met-2* (Andersen and Horvitz, 2007) and *hpl-1* (Couteau et al., 2002) which are expressed within specific neurons (Taylor et al., 2021) showed lower levels of behavioral time-synchronization across multiple developmental stages (L2-Adult), compared to the wild-type population (Fig. 2A-E; Fig. S2A). Similarly, mutants for the *npr-1* neuropeptide receptor which were previously measured for their long-term roaming activity across development (Stern et al., 2017; Harel et al., 2024), showed a significant decrease in the temporal synchronization among individuals during the L3-Adult developmental stages, but a strong increase in behavioral synchronization during the L1 stage (Fig. 2A-C,F; Fig. S2A). In contrast to the identified decrease in the developmental synchronization of behavior, we also found that in a fraction of the mutant populations the behavioral time-synchronization was increased across most of the developmental stages. For example, mutants for the *osm-9* channel involved in sensation of environmental inputs (Colbert et al., 1997) and *tph-1* serotonin-deficient individuals showed significantly higher developmental correlations compared to the wild-type population during most of the stages (Fig. 2A-C,G; Fig. S2B,D). Interestingly, some of the effects on the temporal synchronization were highly limited in time, as *tax-4* mutant individuals that are sensory-defective (Bargmann et al., 1993; Dusenbery et al., 1975; Mori and Ohshima, 1995) showed higher levels of behavioral time-synchronization during the L2 stage (Fig. 2A-C,H; Fig. S2B,D).

**Figure 2.**
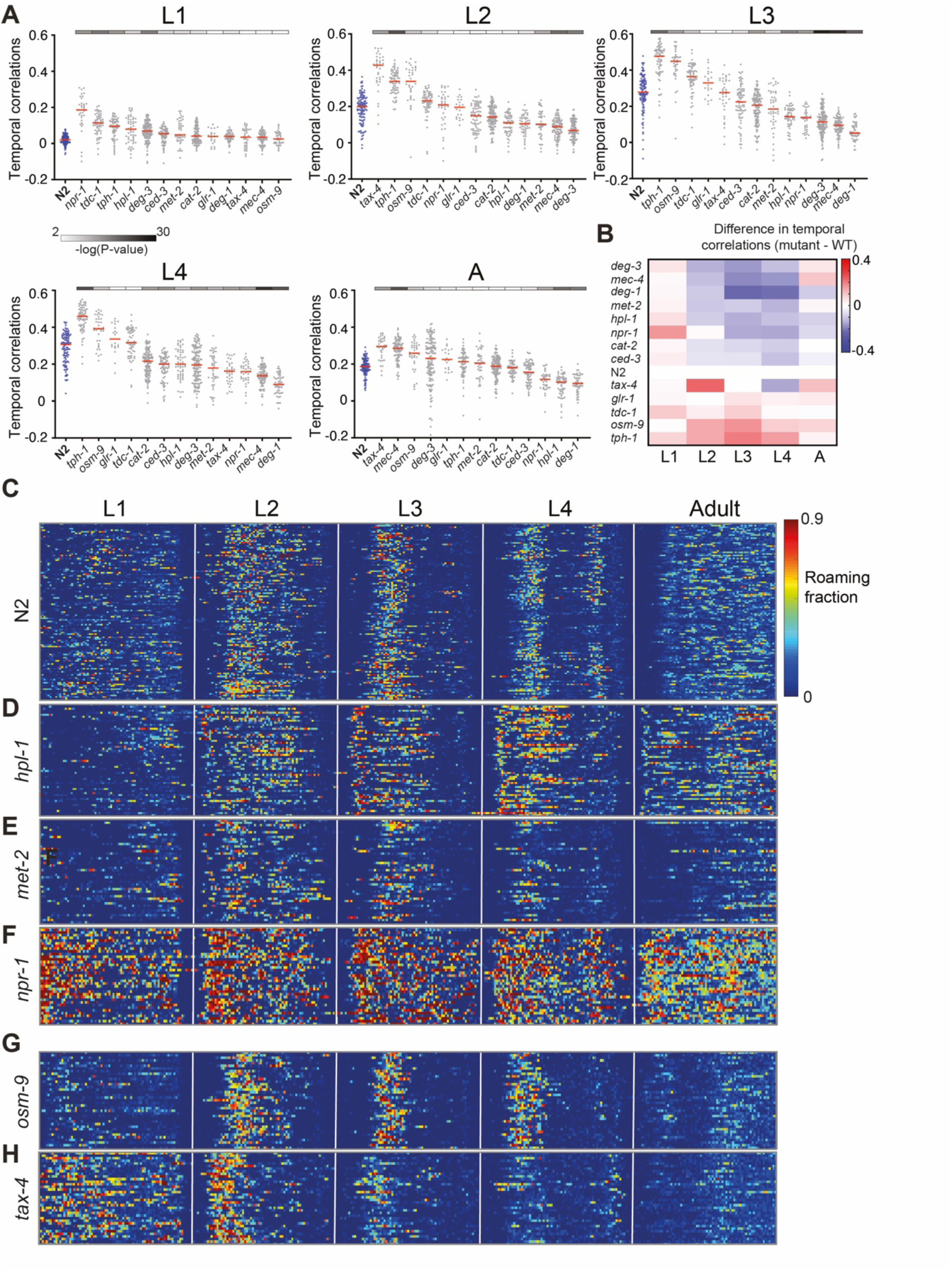
Uncovering neuronal pathways affecting behavioral time-synchronization across development. **(A)** Temporal correlations of roaming behavior among individuals within the wild-type population (blue) and 13 mutant populations for neuronal genes (grey) in each developmental stage. Each dot represents the average temporal correlation between a single individual and all other individuals in the population. Red bars represent the median correlation across each of the populations. Mutant populations are sorted based on their median temporal correlation for visualization. Color bars (top) represent the significance (-log(P-value)) of the difference in temporal correlations between each of the mutant populations and the wild-type (Wilcoxon rank-sum test, FDR corrected). Indicated are P-values<0.01. **(B)** Difference in median temporal correlation between each of the mutant populations and the wild-type in each developmental stage. **(C-H)** Roaming and dwelling behavior of (C) wild-type, and (D) *hpl-1,* (E) *met-2*, (F) *npr-1*, (G) *osm-9*, (H) *tax-4* mutant individuals. Each row indicates the behavior of one individual across all developmental stages. Different stages are separated by white lines indicating the middle of the lethargus state. Color bar represents the fraction of time spent roaming in each of the time-bins. N2 n=113, *deg-1(u38)* n=51, *mec-4(u231)* n=93, *deg-3(u662)* n=139, *hpl-1(n4317)* n=56, *met-2(n4256)* n=44, *npr-1(ad609)* n=37, *osm-9(ky10)* n=37, *tax-4(p678)* n=36, *ced-3(n717)* n=75, *glr-1(n2461)* n=20, *tph-1(mg280)* n=67, *cat-2(e1112)* n=124, *tdc-1(n3420)* n=60.

While we were able to detect strong effects on the developmental synchronization of behavior in a fraction of the mutant populations, other populations showed milder effects on behavioral time-synchronization. In particular, we found that populations mutant for the *cat-2* and *tdc-1* genes required for the synthesis of specific neuromodulators (dopamine and tyramine/octopamine, respectively), as well as individuals deficient for the glutamate receptor *glr-1* and the apoptotic *ced-3* gene in which the number of neurons is increased, all showed lower levels of impairment of the behavioral time synchronization (Fig. 2A-C; Fig. S2C,E-H). Overall, these results identify specific neuronal pathways that affect the robustness of developmental synchronization of behavior across distinct developmental stages.

### Degeneration of mechanosensory neurons impairs the developmental synchronization of behavior

In addition to other sensory modalities, as part of our screen, we also tested the effects of mechanosensory circuits on the developmental synchronization of behavior. In particular, we have analyzed multiple neurodegenerative mutants in which hyperactive forms of the degenerins *deg-1*, *deg-3,* and *mec-4* lead to cell-autonomous degeneration of specific mechanosensory neurons (Chalfie and Wolinsky, 1990; Driscoll and Chalfie, 1991; Treinin and Chalfie, 1995). Surprisingly, we found that all three neurodegenerative mutant populations showed significant impairment of the temporal synchronization of behavior, compared to the wild-type (Fig. 2A-B; Fig. 3; Fig. S3B-D).

**Figure 3.**
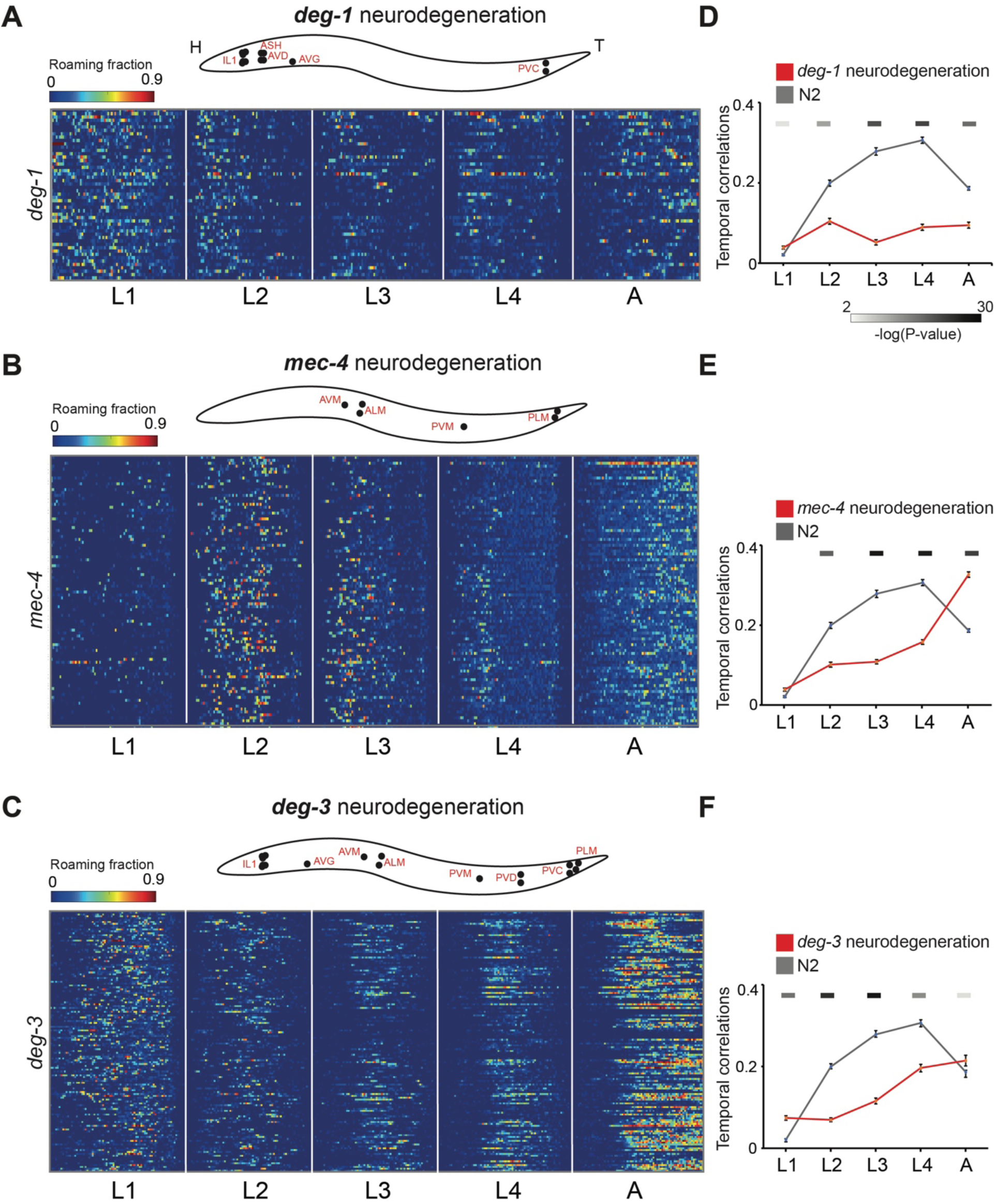
Mechanosensory circuit degeneration impairs the developmental synchronization of long-term behavior. **(A-C)** Illustrations of known degenerated neurons (top) and roaming and dwelling behavior (bottom) in (A) *deg-1,* (B) *mec-4*, and (C) *deg-3* neurodegenerative mutant individuals. Each row indicates the behavior of one individual across all developmental stages. Different stages are separated by white lines indicating the middle of the lethargus state. Color bar represents the fraction of time spent roaming in each of the time-bins. ’H’ and ‘T’ mark head and tail, respectively. **(D-F)** Temporal correlations of roaming behavior among individuals within the (D) *deg-1,* (E) *mec-4*, and (F) *deg-3* mutant populations (red), compared to the wild-type (grey), across distinct developmental stages. Error bars represent standard error of the mean. Color bars (top) represent the significance (-log(P-value)) of the difference in temporal correlations between each of the mutant populations and the wild-type (Wilcoxon rank-sum test, FDR corrected). Indicated are P-values<0.01. N2 n=113, *deg-1(u38)* n=51, *mec-4 (u231)* n=93, *deg-3(u662)* n=139.

In particular, the *deg-1* and *mec-4* genes encode subunits of sodium ion channels involved in mechanosensation (Chalfie and Sulston, 1981; Cueva et al., 2007; Geffeney et al., 2011; O’Hagan et al., 2005; Suzuki et al., 2003). The localization of hyperactive forms of these channels to specific neuronal circuits may generate an imbalance of ion flow (Bianchi et al., 2004), causing cell death by necrosis.

The expression of the *deg-1* hyperactive channel leads to degeneration of a limited set of neurons including ASH, IL1, AVD, AVG, and PVC (Chalfie and Wolinsky, 1990; Hall et al., 1997). Analyses of developmental synchronization of behavior in *deg-1* mutant individuals showed that behavioral time-synchronization throughout development time was strongly impaired, as temporal correlations were decreased during the L2-Adult stages (Fig. 3A,D).

Similarly, in hyperactive mutants for the *mec-4* channel, a highly localized mechanosensory circuit which is composed of only six neurons is degenerated (pair of ALM, pair of PLM, AVM, PVM) (Driscoll and Chalfie, 1991; Suzuki et al., 2003). We found that the developmental synchronization of behavior across larval stages in these neurodegenerative mutants was significantly altered. In particular, similar to the *deg-1* mutants, during the L2-L4 stages *mec-4* neurodegenerative mutants showed lower behavioral time-correlations (Fig. 3B,E), implying that in *mec-4* neurodegenerative mutants the consolidation of developmental synchronization of behavior may be affected during intermediate developmental stages. Furthermore, the analyzed hyperactive mutant forms of *deg-1* and *mec-4* were selected based on their known strong neurotoxic effect, which leads to neurodegeneration early in development (*deg-1(u38)* and *mec-4(u231)*) (Hall et al., 1997). Analysis of an additional *mec-4* hyperactive mutant allele in which neurodegeneration phenotypes appear later during larval stages (*mec-4(e1611))* (Hall et al., 1997), probably due to a milder neurotoxic effect, showed a strong disruption of the normal synchronized activity peaks, but also the generation of alternative short-term peaks (Fig. S3E,F).

In contrast to the degenerins *deg-1* and *mec-4* which encode sodium ion channels, the gene *deg-3* encodes a subunit of a nicotinic acetylcholine receptor (Treinin and Chalfie, 1995). The hyperactive form of *deg-3* leads to similar necrotic degeneration in a larger group of neurons overlapping with those of *deg-1* and *mec-4* (including PLM, PVC, PVD, PVM, ALM, AVM, AVG and IL1). Similar to *deg-1* and *mec-4* neurodegenerative mutants, analysis of *deg-3* mutants revealed lower temporal correlations of behavior during multiple developmental stages (L2-L4 larval stages) (Fig. 3C,F). In summary, these results suggest that the degeneration of specific mechanosensory neurons within the nervous system by different molecular pathways alters developmental synchronization of long-term behavior.

### Early activity patterns in downstream motor neurons are altered following the mechanosensory circuit degeneration

Since the degeneration of mechanosensory neurons disrupts the developmental synchronization of behavior, we aimed to further test potential changes in the underlying neuronal activity dynamics caused by this neurodegeneration. Specifically, we focused on studying alterations in neuronal activity patterns resulting from the degeneration of the six mechanosensory neurons in *mec-4* hyperactive mutants (Driscoll and Chalfie, 1991). We explored the possibility that the degeneration of this relatively small mechanosensory circuit could lead to a broader destabilization of neuronal activity patterns in other downstream circuits. Since ventral nerve cord (VNC) motor circuits in *C. elegans* are shaped post-embryonically and are known to be involved in regulating locomotory patterns (Sulston and Horvitz, 1977; Von Stetina et al., 2005; Mulcahy et al., 2022; Qi et al., 2013), we hypothesized that their function may be affected by the early onset of the mechanosensory circuit degeneration.

To test the effect of the mechanosensory circuit degeneration on the activity patterns of tens of downstream motor neurons we monitored their calcium dynamics in individuals in which the mechanosensory circuit is degenerated, as well as in the wild-type population in which the mechanosensory circuit is intact (Fig. 4A; Fig. S4A). In L4 wild-type individuals, motor neurons showed relatively slow neuronal activity dynamics (average duration of activity bouts: ∼0.85 min). In addition, the temporal patterns of spontaneous activity in different motor neurons are highly correlated within the individual, and usually reflected by at least two groups of neurons that show opposite activity dynamics (Fig. 4B). By quantifying the neuronal activity of motor neurons in L4 individuals in which the mechanosensory circuit is degenerated, we found that activity patterns in these neurons were impaired following the mechanosensory circuit degeneration. In particular, neuronal alterations were reflected by the significantly higher amplitudes of neuronal activity during activity bouts (Fig. 4C) and longer durations of activity bouts in motor neurons of the mechanosensory neurodegenerative individuals (Fig. 4D). In contrast, the frequency of activity bouts over time was lower in the motor neurons, following the degeneration of the mechanosensory circuit (Fig. 4E). Moreover, similar profiles of neuronal destabilization were evident across various types of motor neurons (Fig. S4B). To further ask if the altered activity states of the motor neurons are established early during development, we repeated our measurement in second larval stage (L2) individuals. Interestingly, similar to L4 individuals, we found an increase in activity amplitudes and average duration of activity bouts also in younger L2 individuals in which the mechanosensory circuit is degenerating (Fig. 4F-G; Fig. S4C). However, in contrast to L4 individuals, the frequency of activity bouts following neurodegeneration in L2 individuals was similar to the wild-type population (Fig. 4H), suggesting that specific alterations of neuronal activity states in downstream motor neurons are established later in development. These results suggest that neurodegeneration of a local mechanosensory circuit modifies the underlying neuronal activity dynamics within the nervous system across developmental stages.

**Figure 4.**
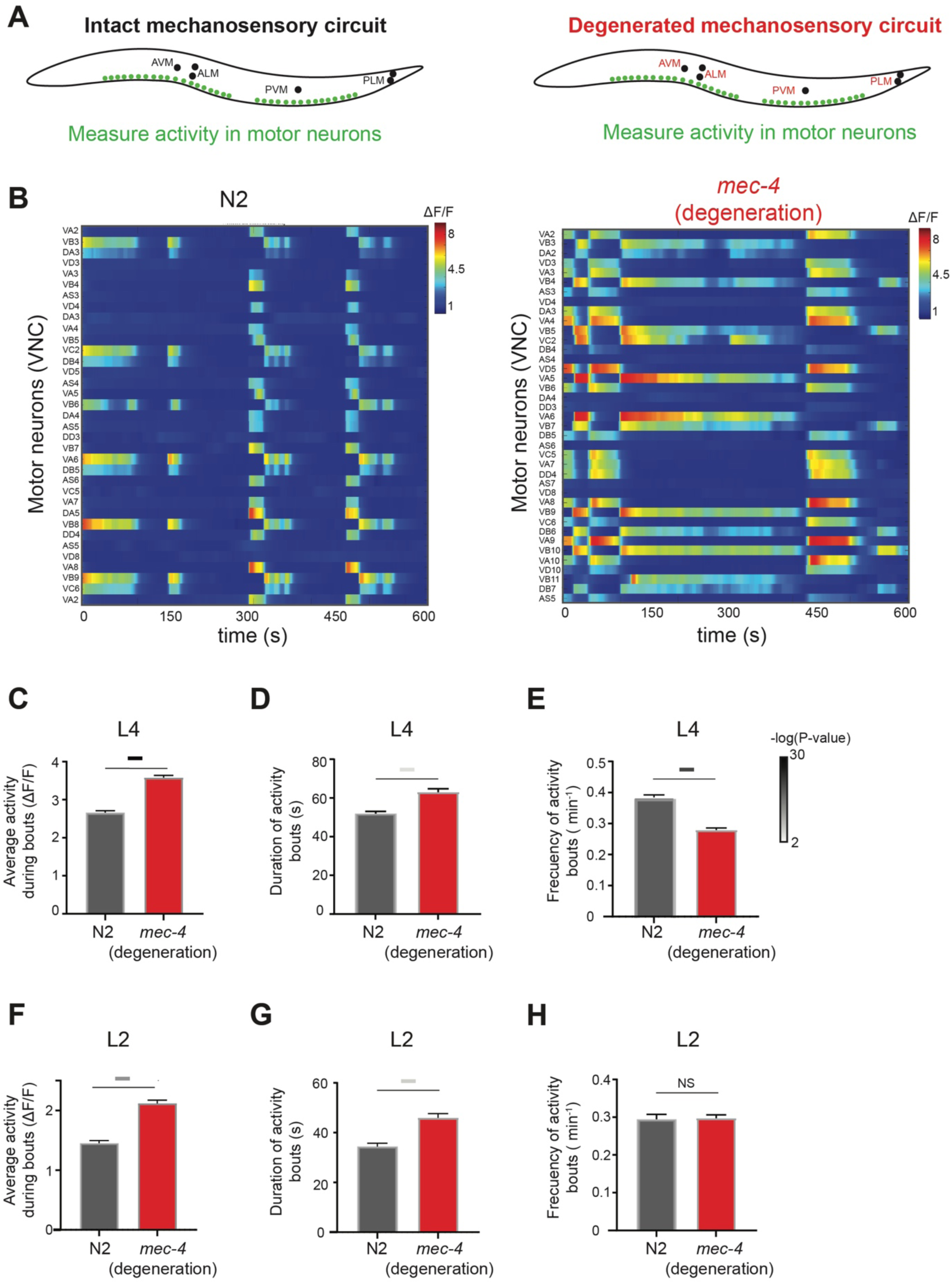
Altered neuronal activity dynamics in downstream motor neurons following the degeneration of a mechanosensory circuit. **(A)** Scheme representing neuronal activity measurements of VNC motor neurons in N2 wild-type individuals (Left) and *mec-4* neurodegenerative mutants *(u231*) in which the touch receptor neurons are degenerated (Right). **(B)** Heat maps represent examples of spontaneous neuronal activity dynamics in motor neurons of wild-type (Left) and *mec-4* neurodegenerative mutant individuals (Right) across 10 minutes in the L4 stage. Color bar represents neuronal activity (ΔF/F), smoothed using moving median (10 bins). **(C-E)** Average neuronal activity during bouts (C), duration of activity bouts (D), and frequency of activity bouts (E) in motor neurons of L4 *mec-4* neurodegenerative mutants (red), compared to wild-type (grey) (see methods). (**F-H)** Average neuronal activity during bouts (F), duration of activity bouts (G), and frequency of activity bouts (H) in motor neurons of L2 *mec-4* neurodegenerative mutants (red), compared to the wild-type (grey). *mec-4(u231)* neurodegenerative mutants (L4: n=615 neurons, 21 individuals; L2: n=351 neurons 15 individuals). N2 (L4: n=722 neurons, 24 individuals; L2: n=185 neurons, 10 individuals).

### Neuroprotection of the mechanosensory circuit from degeneration restores behavioral synchronization with development time

The requirement of the mechanosensory circuit for establishing normal behavioral synchronization across development led us to ask whether neuroprotection from degeneration of the mechanosensory circuit would reestablish normal modes of developmental synchronization of behavior (Fig. 5A). The function of the mechanosensory neurons was shown to be dependent on the *mec-4* channel itself only during specific mechanosensory contexts (Suzuki et al., 2003). Thus, we initially tested whether the degeneration of *mec-4* expressing neurons in the mechanosensory circuit is required for the impairment of synchronized behavioral structures, or whether the loss of function of the *mec-4* channel itself within these neurons is sufficient to reproduce the same effect. To study this, we measured the behavior of two loss-of-function mutant alleles of the *mec-4* gene which impair the function of the channel within the neurons without causing neurodegeneration of the mechanosensory circuit. We found that in contrast to the effect of the mechanosensory circuit degeneration, the loss-of-function *mec-4* mutants did not show the decrease in the temporal synchronization of behavioral patterns during development (Fig. 5B,E; Fig. S5A-D,G-H). In addition, chemogenetic inhibition of the circuit by the cell-specific expression of HisCl (Pokala et al., 2014) showed only minor effects on the temporal behavioral patterns (Fig. S5K). These results suggest that the degeneration of the mechanosensory circuit is required for the alteration of the normal developmental synchronization of behavior.

**Figure 5.**
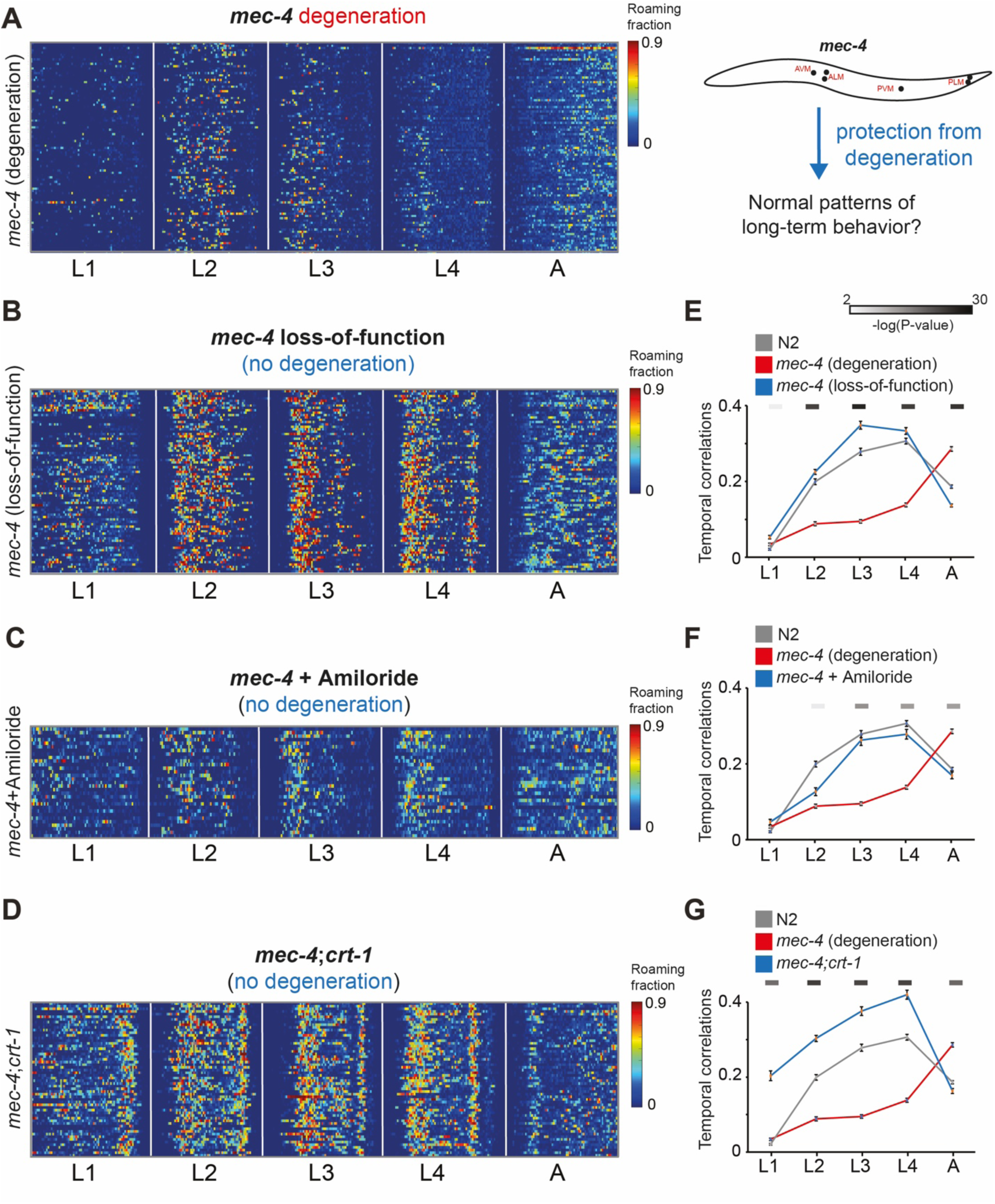
Neuroprotection from degeneration reestablishes normal patterns of developmentally synchronized behaviors. **(A)** Left: Roaming and dwelling behavior in *mec-4* neurodegenerative mutant individuals as in 3B. Each row indicates the behavior of one individual across all developmental stages. Different stages are separated by white lines indicating the middle of the lethargus state. Color bar represents the fraction of time spent roaming in each of the time-bins. Right: Illustration of protection from *mec-4* degeneration. **(B-D)** Roaming and dwelling behavior in (B) *mec-4* loss-of-function mutants *(e1339)* not causing degeneration, (C) *mec-4* neurodegenerative mutants *(u231)* with the supplementation of amiloride, and (D) double mutants for the *mec-4* neurodegenerative mutation *(u231)* and *crt-1(bz29)* mutation. **(E-G)** Temporal correlations of roaming behavior among individuals within the (E) *mec-4* loss-of-function mutants *(e1339)* not causing degeneration, (F) *mec-4* neurodegenerative mutants *(u231)* with the supplementation of amiloride, and (G) double mutants for the *mec-4* neurodegenerative mutation (u231) and *crt-1 (bz29)* mutation (blue), compared to the wild-type (grey) and *mec-4* neurodegenerative mutant individuals *(u231)* (red), across distinct developmental stages. Error bars represent standard error of the mean. Color bars (top) represent the significance (-log(P-value)) of the difference in temporal correlations between each of the neuroprotected populations and the *mec-4* neurodegenerative mutants *(u231)* (Wilcoxon rank-sum test, FDR corrected). Indicated are P-values<0.01. N2 n=113, *mec-4(u231)* n=93, *mec-4(e1339)* n=93, *mec-4(u231)* + amiloride n=31, *mec-4(u231)*;*crt-1(bz29)* n=58.

Next, we examined whether preventing the mechanosensory circuit degeneration by using the drug amiloride, a known blocker of degenerin sodium channels (Bianchi et al., 2004; Brown et al., 2007; Goodman et al., 2002), would restore the normal behavioral synchronization across development. We found that supplementing exogenous amiloride to the *mec-4* neurodegenerative mutant individuals which normally show the impairment of developmental synchronization of behavior, led to the reestablishment of synchronized behaviors across developmental stages (Fig. 5C,F; Fig. S5E,I). These results imply that protection from mechanosensory circuit degeneration by an external drug restores the normal developmental synchronization of behavior. Furthermore, it has been previously shown that mutation of the calmodulin *crt-1* which is involved in homeostasis of cellular calcium reservoir from the ER, genetically suppresses the neurotoxic effect in *mec-4* neurodegenerative mutants (Xu et al., 2001). In contrast to the *mec-4* limited expression in the nervous system, *crt-1* is broadly expressed across many neurons (Park et al., 2001; Taylor et al., 2021) and *crt-1* single mutants showed additional synchronized roaming activity peaks at the end of each larval stage (Fig. S5J). We found that in *crt-1;mec-4* double mutants in which the neurodegeneration of the mechanosensory circuit is prevented, the impairment of behavioral synchronization is eliminated (Fig. 5D,G; Fig. S5F). Overall, these results show that pharmacological or genetic neuroprotection of the mechanosensory circuit from neurodegeneration restores normal modes of developmentally-synchronized behaviors.

## Discussion

Across different species, long-term behavioral patterns are highly structured across developmental timescales, reflected in predicted changes in behavioral outputs that depend on the specific developmental window. It has previously been shown that during development, *C. elegans* individuals exhibit temporal patterns of long-term behavior that are highly dynamic across and within developmental stages, shaped by both the neuromodulatory state of the population and its environmental experiences (Stern et al., 2017; Harel et al., 2024; Ali Nasser et al., 2023). In this work, we studied how developmental synchronization of behavior is established across the entire developmental trajectory. In addition, we studied the neuronal circuits within the nervous system that are required for this robust developmental synchronization. Our results revealed that wild-type individuals showed variable levels of temporal synchronization of behavior across distinct developmental stages. Interestingly, while individuals exhibited near-random temporal patterns of behavior during the first larval stage, synchronization of behavioral patterns with development time gradually increased during later larval stages. These results suggest the consolidation of developmental patterns of behavior across stages, as development progresses. As the *C. elegans* nervous system is being shaped during post-embryonic development, both at the cell state (Ripoll-Sánchez et al., 2023; Sun and Hobert, 2021; Taylor et al., 2021) and structural level (Sulston and Horvitz, 1977; White et al., 1986; Witvliet et al., 2021) to affect behavior, the increase in the robustness of time-locked behavioral patterns across developmental stages may reflect the control of developmentally-synchronized behaviors by the function of specific circuits that are shaped post-embryonically.

Following this hypothesis, we sought to identify neuronal pathways that may impair the temporal structure of behavior and its consolidation across developmental stages. By analyzing the developmental synchronization of behavior in multiple mutant populations, where the function or structure of the nervous system is perturbed, we found multiple instances in which the degree of behavioral synchronization across specific developmental stages is either decreased or increased. For example, mutants for the epigenetic chromatin regulators *hpl-1* and *met-2* that are known to be expressed within the *C. elegans* nervous system (Taylor et al., 2021) showed lower developmental synchronization compared to the wild-type, indicating their relatively unstructured behavior over time. Epigenetic mechanisms were shown to act within neurons to control developmental timing of cell maturation (Ciceri et al., 2024). It is plausible that epigenetic alterations within specific circuits may destabilize cellular states required for establishing developmentally synchronized behavioral patterns. In contrast, we also found cases in which long-term behavioral patterns across individuals became more developmentally-synchronized within specific mutant populations. For instance, mutants for *osm-9* or *tax-4* that are known to be involved in multiple sensory modalities (Bargmann et al., 1993; Colbert et al., 1997; Dusenbery et al., 1975; Mori and Ohshima, 1995) showed a higher degree of developmental synchronization of behavior, across most of the developmental stages or during specific stages, respectively. The increase in temporal synchronization in these sensory mutants implies that environmental inputs may also play a role in tuning developmental synchronization of behavior. These results suggest that multiple neuronal pathways can modify the levels of robustness of developmental structures of behavior, further emphasizing the active control of time-locked behavioral patterns across development by the *C. elegans* nervous system.

In addition to gene mutants that affect multiple neuronal functions, we also tested neurodegenerative mutants in which hyperactive forms of *mec-4*, *deg-1* and *deg-3* lead to neurodegeneration of specific mechanosensory neurons relatively early in development (Chalfie and Wolinsky, 1990; Driscoll and Chalfie, 1991; Treinin and Chalfie, 1995). We found that all three degeneration processes, despite being generated via different mechanisms, showed strong impairment of the normal developmental synchronization of behavior. In a particular case in which a hyperactive form of *mec-4* leads to neurodegeneration of only six mechanosensory neurons we further found that following the mechanosensory circuit degeneration, the neuronal activity patterns of tens of downstream motor neurons was destabilized, reflected by the increase in neuronal activity amplitudes and the longer duration of neuronal activity bouts. Interestingly, we have observed these modifications in neuronal activity patterns early in development, suggesting that the early local degeneration of the mechanosensory circuit leads to the alterations of downstram circuits within the nervous system.

We further asked if the neuroprotection of the mechanosensory circuit could reverse the long-term alterations of behavioral synchronization across development. Our results revealed that either pharmacological or genetic neuroprotection from degeneration restored normal developmental synchronization of behavior. These results imply that even within the relatively compact nervous system of *C. elegans*, the degeneration of specific mechanosensory circuits alters the normal developmental patterns of behavior. A plausible hypothesis for the involvement of a mechanosensory circuit in the developmental synchronization of behavior may be its requirement for assessing body size growth across development through mechanosensory inputs from its surface.

More broadly, these results suggest that while under the normal context of an intact nervous system, individuals show highly stereotyped temporal patterns of behavior, alterations of specific modulatory circuits may generate developmental diversity of long-term behavioral patterns among individuals. We hypothesize that on evolutionary timescales, this exposed temporal diversity in behavior within populations may serve as phenotypic parameter that may drive modifications in the nervous system towards alternative patterns of behaviors that are established across the developmental trajectory.

## Methods

### C. elegans populations

*C. elegans* strains used in this study: Wild-type Bristol N2; ZB1028 *crt-1(bz29)* V; TU1948 *deg-1(u38)* X; TU1747 *deg-3(u662)* V; *SST246 mec-4(u231)* X*; crt-1(bz29)* V (*mec-4*::GFP crossed out from ZB1602); CB1611 *mec-4(1611)* X; CB1339 *mec-4(e1339)* X; TU231 *mec-4(u231)* X; TU253 *mec-4(u253)* X; MT15434 *tph-1(mg280)* II; MT1522 *ced-3(n717)* IV; KP4 *glr-1(n2461)* III; MT13971 *hpl-1(n4317)* X; MT13293 *met-2(n4256)* III; DA609 *npr-1(ad609)* X; CX10 *osm-9(ky10)* IV; PR678 *tax4*(p678) III; MT10661 *tdc-1(n3420)* II; CB1112 *cat-2(e1112)* II; HisCl strain: SST243 *stsEx*044 [*mec-4p::HisCl1::SL2:GFP* (50 ng/μl)*; Pmyo-2::mCherry::unc-54utr* (5 ng/μl)]. Neuronal imaging strains: OH16230 *otIs670[NeuroPAL], otIs672[panneuronal::GCaMP6s];* SST116 *mec-4(u231); otIs670[NeuroPAL]; otIs672[panneuronal::GCaMP6s]* was generated by crossing TU231 strain with OH16230 strain.

### Plasmids

*pLL013* (*mec-4p::HisCl1::SL2:GFP*) was generated by replacing the promoter in *pNP472* (Pokala et al., 2014) with 1069 bp of *mec-4* promoter amplified from *Pmec-4::lin-14* plasmid (Ritchie et al., 2017).

### Growth conditions

*C. elegans* worms were maintained on NGM agar plates, supplemented with E. coli OP50 bacteria as a food source. For behavioral tracking, we imaged single individuals grown from egg-hatching to adulthood in custom-made laser-cut multi-well plates. Each well (10mm in diameter) was seeded with a specified amount of OP50 bacteria (10 µL of 1.5 OD) that was UV-killed before the experiment to prevent bacterial growth during the experiment. For the amiloride supplementation experiments, 30µL of amiloride (10mM) (Sigma, A7410) was applied to each well in the imaging plates after bacterial seeding and UV illumination. In addition, the source plates used for embryos extraction contained OP50 with a concentration of 10mM amiloride. For the HisCl chemogenetic experiments (Pokala et al., 2014), 10µl of Histamine (200mM) (Sigma, H7250) was applied to each well in the imaging plates after bacterial seeding and UV illumination.

### Behavioral imaging system

Longitudinal behavioral imaging was performed using custom-made imaging systems. Each imaging system consists of an array of six 12 MP USB3 cameras (FLIR, Flea3) and 35 mm high-resolution objectives (Edmund optics) mounted on optical construction rails (Thorlabs). Each camera images six individual wells. Movies are captured at 3 fps with a spatial resolution of ∼9.5 um. For homogenous illumination of the imaging plates we used identical LED backlights (Metaphase Technologies) and polarization sheets. Imaging was conducted inside a custom-made environmental chamber in which temperature was controlled using a Peltier element (TE technologies, temperature fluctuations in the range of 22.5 ± 0.1°C). Humidity was held in the range of 50% +/− 5% with a sterile water reservoir and outside illumination was blocked, keeping the internal LED backlights as the only illumination source. Movies from the cameras were captured using commercial software (FlyCapture, FLIR) and saved on two computers (3 cameras per computer).

### Extraction of developmental trajectories of behavior

To extract behavioral trajectories of each individual’s center of mass across the experiment, captured videos were analyzed by a custom-made script programmed in MATLAB (Mathworks) (Stern et al. 2017). In each video frame, each individual worm is automatically detected as a moving object in space by background subtraction, and the coordinates of its visual center of mass are logged. In each experiment, 600,000-700,000 frames per individual are analyzed using ∼50 processor cores to reconstruct the full behavioral trajectory over days of measurements across all stages of development. The total time of image processing was 3-4 days for tens of individuals per experiment across the full developmental trajectory. Egg hatching time of each individual is automatically marked by the time when activity can be detected in the arena. The middle of the lethargus periods were manually identified as the transition points between different stages of development (based on 10s-time-scale speed trajectories, smoothed over 300 frames). To align the temporal behavioral trajectories of different individuals we age-normalized individuals by dividing the behavioral trajectory of each developmental stage into a fixed number of time windows.

### Quantification of behavioral parameters

For each individual, we differentiate between roaming and dwelling locomotory states by averaging speed (μm/s) and angular velocity (absolute deg/s) over 10 seconds using a moving time window, and generating a 2D probability distribution of these two behavioral parameters for all intervals in each time-bin along the experiment (50 X 50 bins distribution, speed bin size: 7.59 um/s, angular velocity bin size: 3.6 deg/s) (Stern et al., 2017). Drawing a diagonal through the probability distribution separated roaming and dwelling states (Arous et al., 2009; Flavell et al., 2013; Stern et al., 2017). The behavior of each animal across time was quantified as a sequence of roaming and dwelling intervals. The fraction of time spent roaming represents the fraction of these intervals classified as roaming states within a given time bin. For each developmental stage, we quantified the two-dimensional probability distribution of the whole population and changed the slope of the diagonal to classify roaming and dwelling appropriately (slopes: 5,2.5,2.3,2,1.5 for the L1,L2,L3,L4 and adult stages, respectively). To analyze the temporal synchronization in roaming behavior we quantified the Pearson correlation among each pair of individuals (pairwise temporal correlations) within the population. For comparing the temporal correlations between populations and across developmental stages, the average temporal correlation between each individual and all other individuals in the population was quantified.

### Microscopy and neuronal imaging

For neuronal GCaMP imaging, individual NeuroPAL worms (Yemini et al., 2021) were transferred to a non-seeded NGM plate for 5 minutes to remove residual bacteria, then mounted on a slide with a 2% agar pad and immobilized using 5 mM tetramisole in M9 buffer.

Ventral nerve cord motor neurons were imaged using a Nikon Eclipse Ti2 inverted microscope and 40X objective (NA 0.95). Initial images for neuronal positioning in individuals were captured for pan-neuronal RFP fluorescence. For L4 individuals, imaging neuronal activity was performed by tracking GCaMP fluorescence at 1 Hz for 10 minutes. For L2 individuals, imaging neuronal activity was performed at 0.5 Hz for 15 minutes. Images were taken following the RVG site to the posterior end. The number of VNC motor neurons that were captured in each individual varied due to worm’s position on the slide.

Neuronal activity analysis was performed using Fiji software. Neuronal positions were manually marked using the acquired pan-neuronal RFP images. ΔF/F_0_ was calculated for each neuron (F_0_ defined as the 10th percentile) and data were smoothed using a moving median (10 bins). Neuronal activity thresholds were determined using the Otsu method via MATLAB’s ‘multithresh’ function as previously described (Ji et al., 2021). Identification of VNC neurons was performed manually in L4 individuals based on the neuronal position map (Skuhersky et al., 2022). Identification of single neurons was not performed in L2 individuals due to larger positional variation. Activity bouts were defined as contiguous periods where ΔF/F_0_ values remained above the threshold, bounded by values below the threshold. Based on the identified activity bouts across neurons, averages of activity intensity during bouts (ΔF/F_0_), bouts duration, and bouts frequency were quantified

## Acknowledgments

We thank Monica Driscoll, Cori Bargmann, and Massimo Hilliard for sharing strains and plasmids. We thank members of our laboratory for discussions and comments on the manuscript. Some strains were provided by the CGC, which is funded by the NIH Office of Research Infrastructure Programs (P40 OD010440). This work was supported by the European Research Council ERC-2019-STG and the Israel Science Foundation grant 3035/20.

---

**Figure S1.**
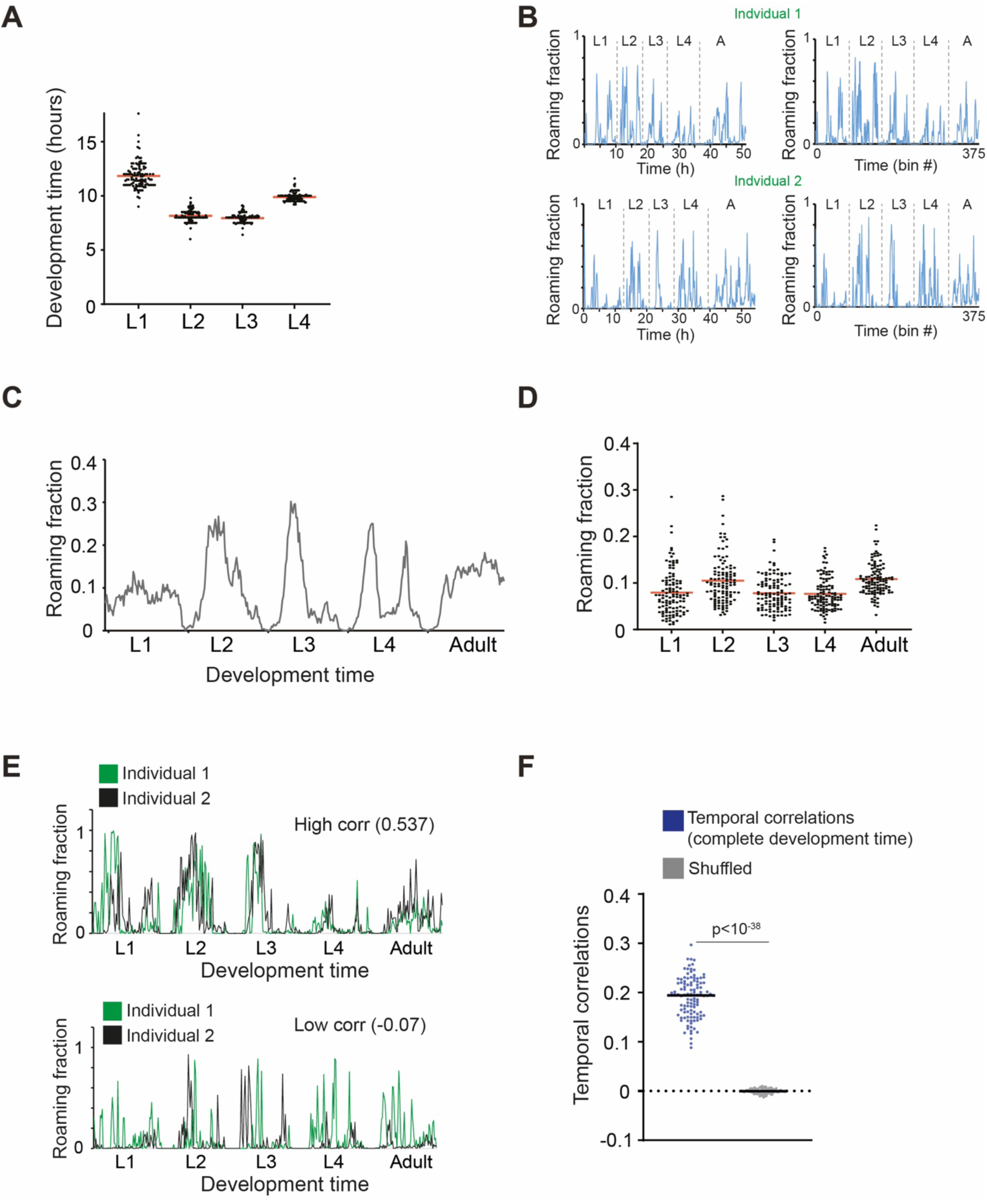
Quantification of development-time, average roaming activity and long-term behavioral synchronization across wild-type individuals. **(A)** Development time across L1-L4 larval stages within the wild-type N2 population (n=113). Each dot represents a single individual. Red bars mark the average across the population. **(B)** Examples of absolute time (left) and age-normalized (right) roaming of two individuals across development. Normalization aligns behavioral trajectories of different individuals across development by dividing each developmental stage of each individual into 75 time-bins. **(C)** Average roaming fraction of wild-type animals across the developmental trajectory (375 time-bins). **(D)** Average roaming fraction of the wild-type population quantified in each developmental stage. Each dot represents a single individual. Red bars mark the average across the population. **(E)** Examples of pairs of wild-type individuals showing relatively high (top) or low (bottom) temporal correlation between their developmental patterns of roaming behavior across a full development time. **(F)** Temporal correlations of roaming behavior among all individuals within the wild-type population (blue) compared to a randomly shuffled dataset (grey), across the complete developmental trajectory. Each dot represents the average temporal correlation between a single individual and all other individuals in the population. Black bar represents the median correlation within the population. P-value calculated by Wilcoxon rank-sum test.

**Figure S2.**
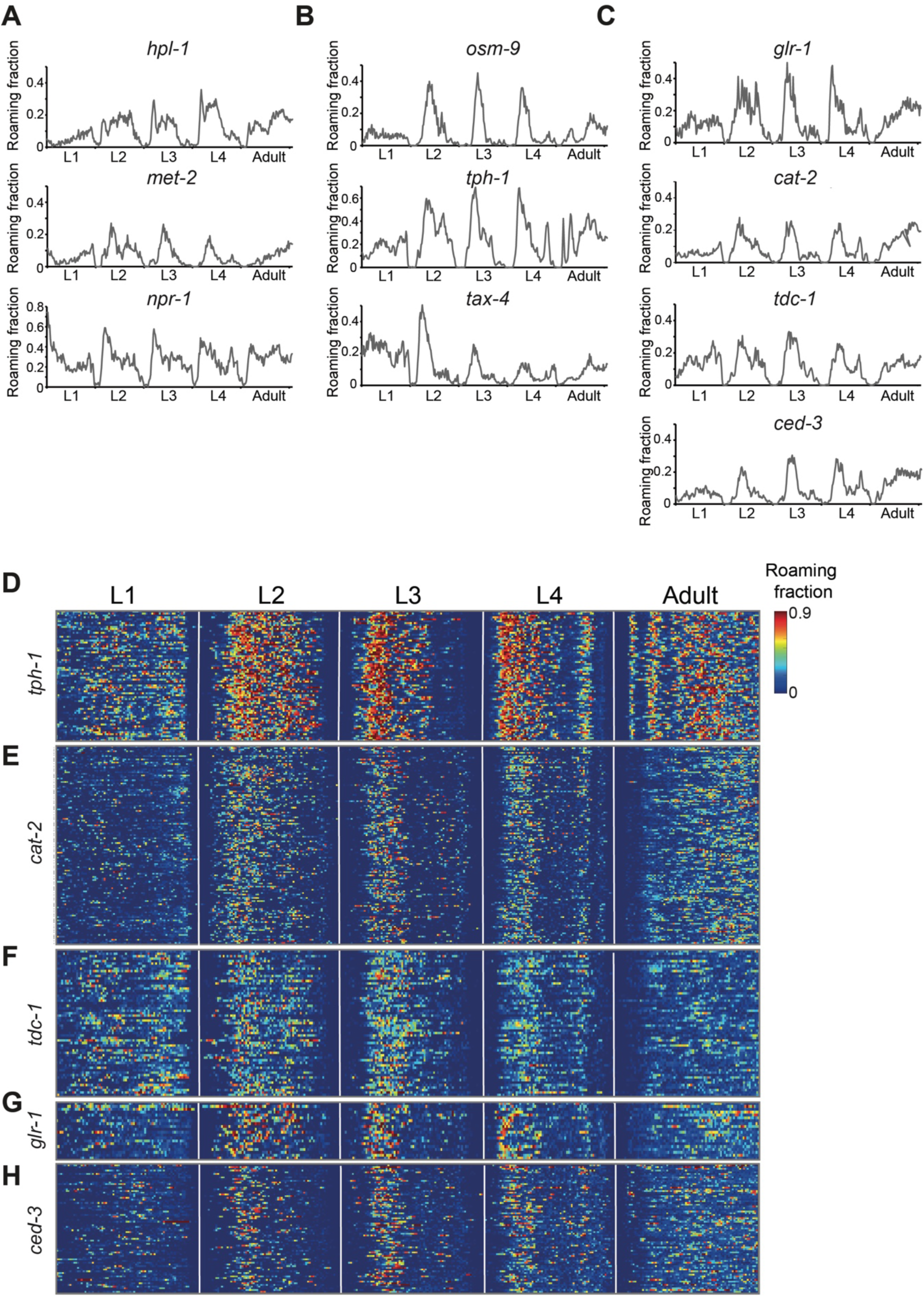
Temporal patterns of roaming activity in neuronal genes mutants. **(A-C)** Average roaming fraction of (A) *hpl-1*, *met-2, npr-1* (B) *osm-9*, *tph-1*, *tax-4* and (C) *glr-1*, *cat-2*, *tdc-1*, *ced-3* mutant individuals across the developmental trajectory. **(D-H)** Roaming and dwelling behavior in (D) *tph-1*, (E) *cat-2*, (F) *tdc-1*, (G) *glr-1*, and (H) *ced-3* mutant individuals. Each row indicates the behavior of an individual across all developmental stages. Different stages are separated by white lines indicating the middle of the lethargus state. Color bar represents the fraction of time spent roaming in each of the time-bins. *tph-1(mg280)* n=67, *cat-2(e1112)* n=124, *tdc-1(n3420)* n=60; *hpl-1(n4317)* n=56, *met-2(n4256)* n=44, *npr-1(ad609)* n=37, *osm-9(ky10)* n=37, *tax-4(p678)* n=36, *glr-1(n2461)* n=20*; ced-3(n717)* n=75.

**Figure S3.**
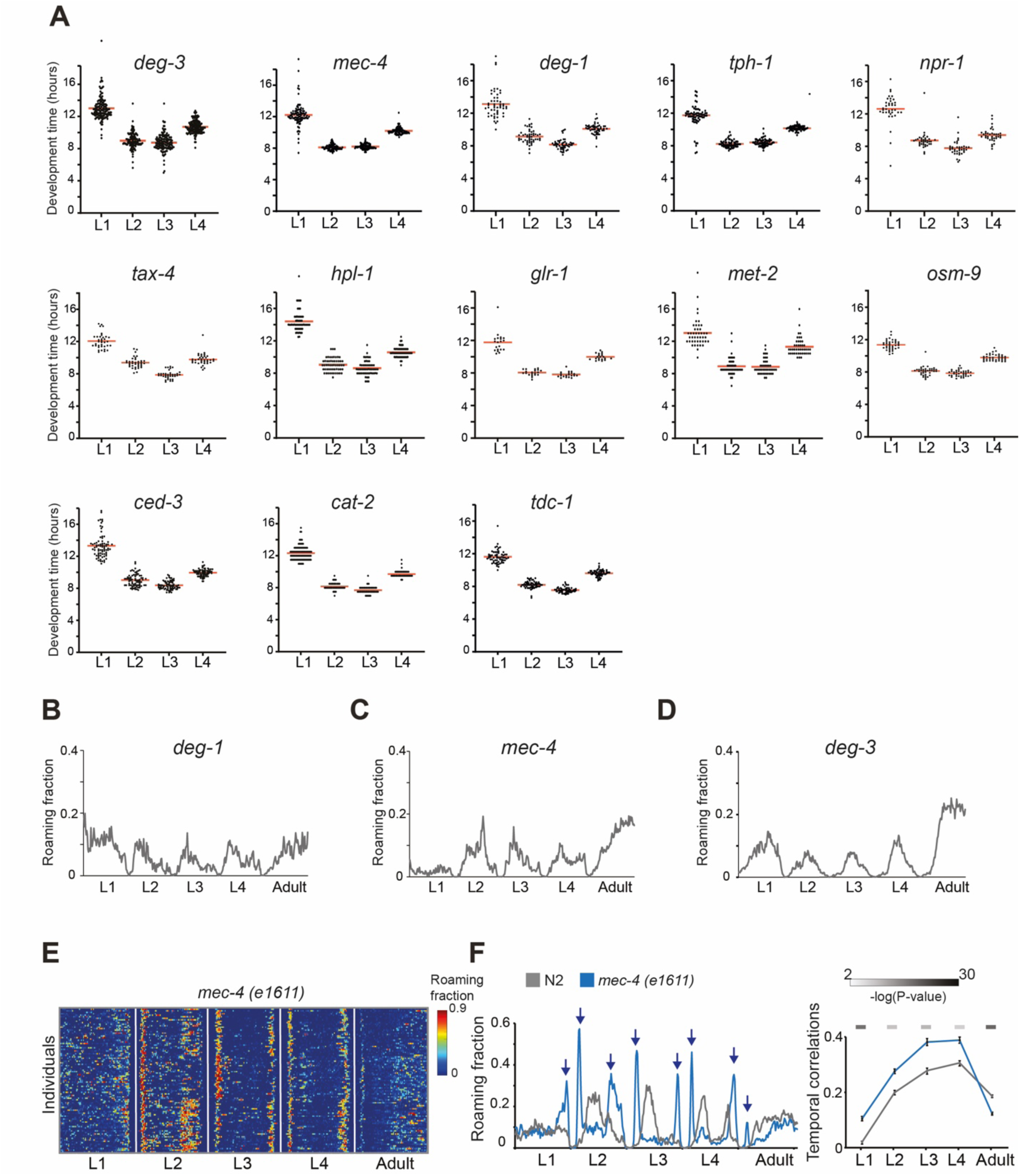
Development time in neuronal genes mutants and alterations in long-term patterns of roaming behavior by neurodegeneration. **(A)** Development time across L1-L4 larval stages within the 13 neuronal genes mutant populations. Each dot represents a single individual. Red bars mark the average across the population. **(B-D)** Average roaming fraction of (B) *deg-1,* (B) *mec-4,* and (C) *deg-3* neurodegenerative mutants across the developmental trajectory. **(E)** Roaming and dwelling behavior in *mec-4(e1611)* individuals. Each row indicates the behavior of one individual across all developmental stages. Different stages are separated by white lines indicating the middle of the lethargus state. Color bar represents the fraction of time spent roaming in each of the time-bins. **(F)** Left: Average roaming fraction of *mec-4 (e1611)* individuals across the developmental trajectory. Arrows mark new synchronized peaks of activity in *mec-4(e1611)* mutant individuals (blue), compared to the wild-type population (grey). Right: temporal correlations of roaming behavior in *mec-4(e1611)* individuals, compared to the wild-type population, across distinct developmental stages. Error bars represent standard error of the mean. Color bars (top) represent the significance (-log(P-value)) of the difference in temporal correlations between the mutant population and the wild-type (Wilcoxon rank-sum test, FDR corrected). Indicated are P-values<0.01. N2 n=113, *deg-1(u38)* n=51, *mec-4(u231)* n=93, *deg-3(u662)* n=139, *mec-4(e1611)* n=99, *tph-1(mg280)* n=67, *cat-2(e1112)* n=124, *tdc-1(n3420)* n=60, *hpl-1(n4317)* n=56, *met-2(n4256)* n=44, *npr-1(ad609)* n=37, *osm-9(ky10)* n=37, *tax-4(p678)* n=36, *glr-1(n2461)* n=20*, ced-3(n717)* n=75.

**Figure S4.**
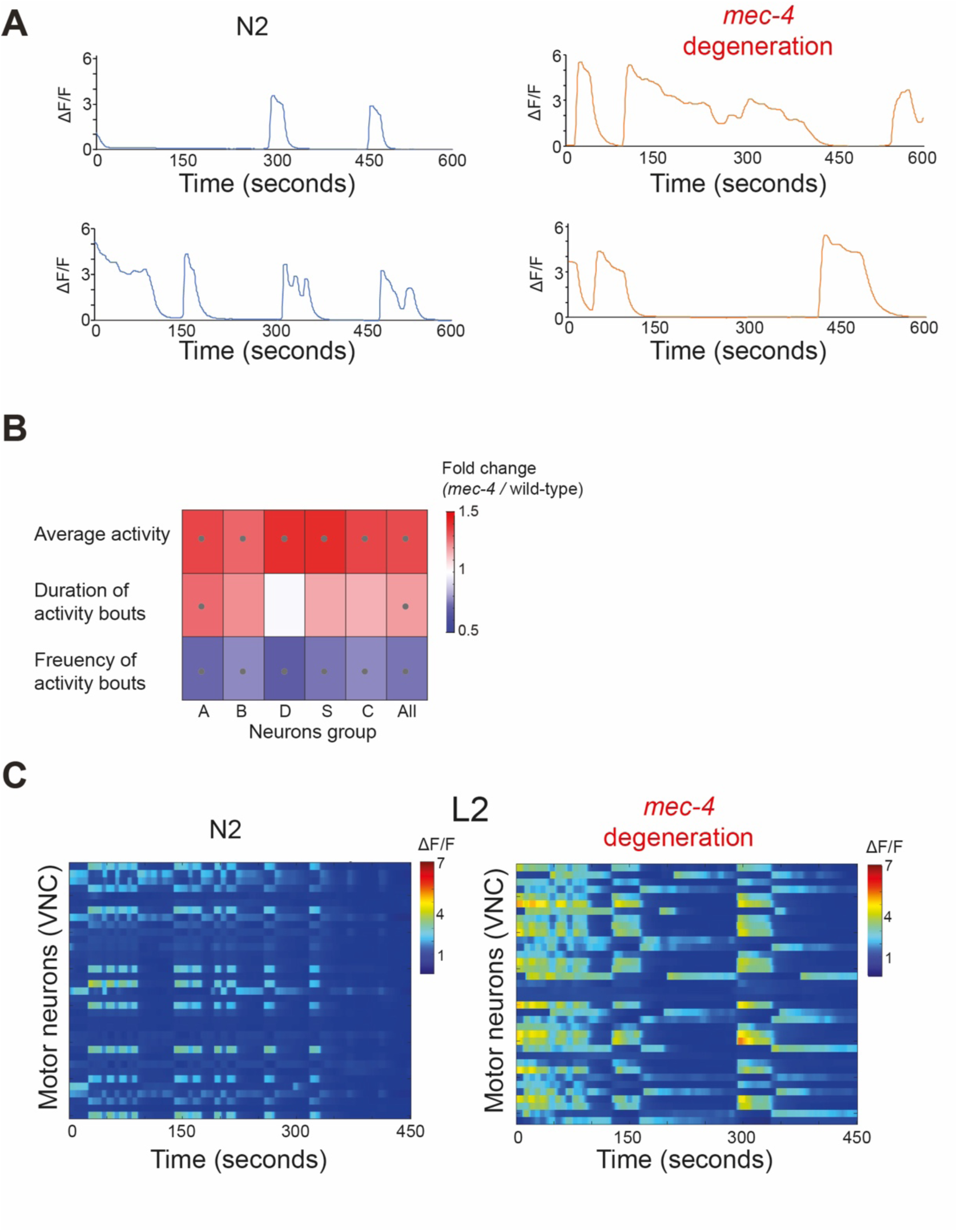
Neuronal activity patterns in motor neurons in wild-type and TRN neurodegenerative mutants. **(A)** Examples of neuronal activity measurements over 10 minutes in single VNC motor neurons in L4 wild-type individuals (left) and *mec-4* neurodegenerative mutants *(u231)* in which the touch receptor neurons are degenerated (right) **(B)** Heatmap represents fold change difference in neuronal activity parameters of different groups of motor neurons between *mec-4* neurodegenerative mutants and wild-type individuals. Grey circles indicate P-value<0.05 (FDR corrected). N2: n=182,184,63,61,36 and *mec-4*: n=144,145,49,53,51 for the A,B,D,S,C motor neuronal groups, respectively. **(C)** Heat maps represent spontaneous neuronal activity dynamics in motor neurons of a wild-type (left) and *mec-4* neurodegenerative mutant individual (right) over 15 minutes in the L2 stage. Color bar represents neuronal activity (ΔF/F), smoothed using moving median (10 bins).

**Figure S5.**
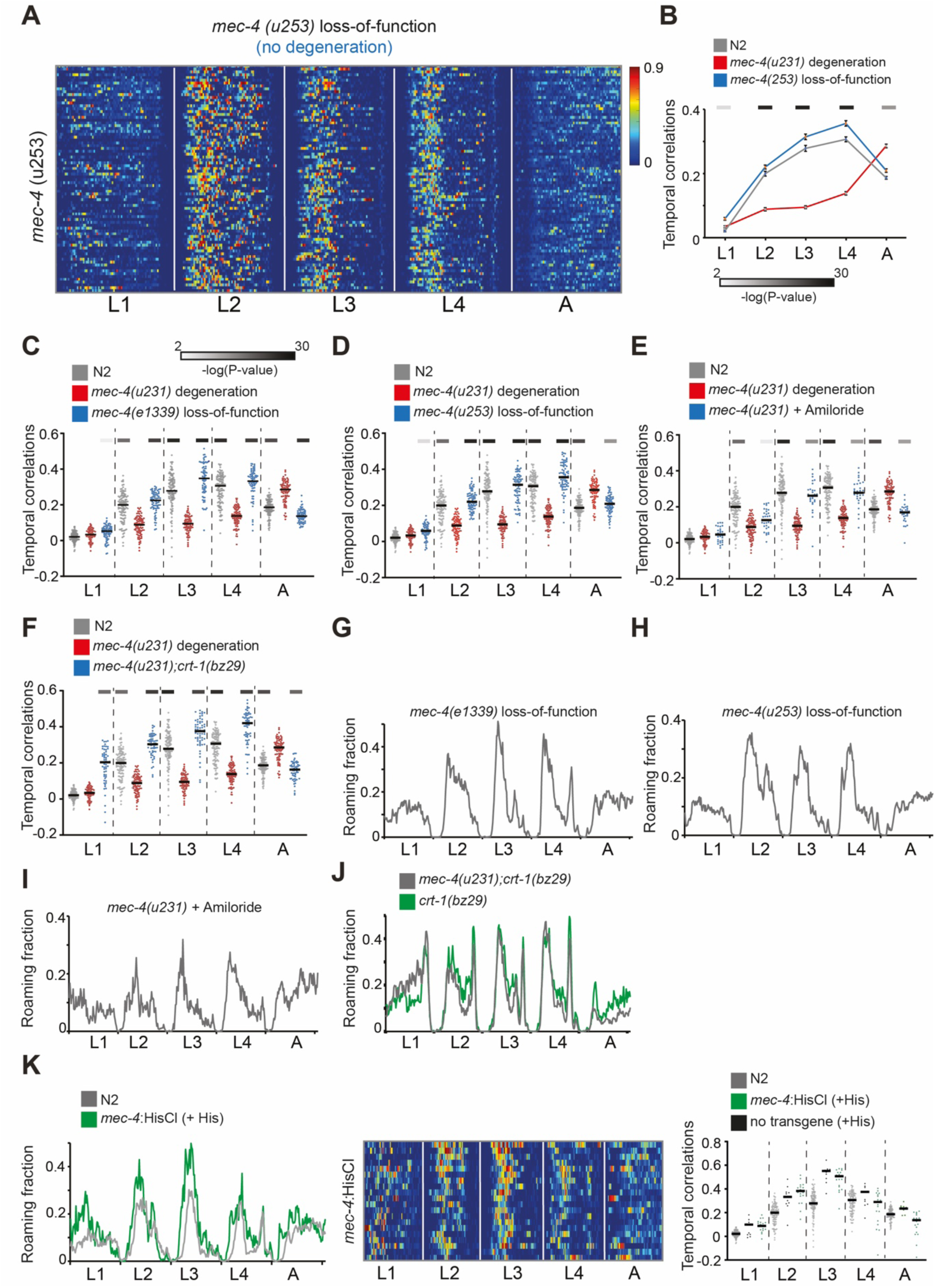
Developmental patterns of behavior following protection from neurodegeneration. **(A)** Roaming and dwelling behavior in *mec-4* loss-of-function *(u253)* mutant individuals. Each row indicates the behavior of one individual across all developmental stages. Different stages are separated by white lines indicating the middle of the lethargus state. Color bar represents the fraction of time spent roaming in each of the time-bins. **(B)** Temporal correlations of roaming behavior among individuals within the *mec-4* loss-of-function *(u253)* mutants (blue), compared to wild-type (grey) and *mec-4 (u231)* neurodegenerative mutant individuals (red), across distinct developmental stages. Error bars represent standard error of the mean. **(C-F)** Temporal correlations of roaming behavior among individuals within the (C,D) *mec-4* loss-of-function mutants (*e1339* and *u253*), (E) *mec-4* neurodegenerative mutants (*u231*) with the supplementation of amiloride, and (F) double mutants for the *mec-4* neurodegenerative mutation (*u231*) and *crt-1* mutation (*bz29*) (blue), compared to the wild-type (grey) and *mec-4* neurodegenerative mutant (*u231*) individuals (red), across distinct developmental stages. Each dot represents the average temporal correlation between a single individual and all other individuals in the population. Black bars represent the median across each of the populations. **(G-J)** Average roaming fraction of (G,H) *mec-4* loss-of-function mutants (*e1339* and *u253*), (I) *mec-4* neurodegenerative mutants (*u231*) with the supplementation of amiloride, (J) double mutants for the *mec-4* neurodegenerative mutation (*u231*) and *crt-1* mutation (*bz29*) (grey), and single mutants of *crt-1* (*bz29*) (green). **(K)** Left: Average roaming fraction of *mec-4*:HisCl individuals, compared to the wild-type population. Middle: Roaming and dwelling behavior of *mec-4*:HisCl individuals. Right: Temporal correlations of roaming behavior among *mec-4*:HisCl individuals, compared to wild-type individuals and individuals that are exposed to histamine and lack the transgene. Color bars in (B-F) represent the significance (-log(P-value)) of the difference in temporal correlations between each of the neuroprotected populations and the *mec-4* neurodegenerative mutants (*u231*) (Wilcoxon rank-sum test, FDR corrected). Indicated are P-values<0.01. *mec-4(u253)* n=80; *mec-4* (e1339) n=93; *mec-4 (u231)* n=93; *mec-4* (u231) + amiloride n=31, *mec-4* (u231);*crt-1*(bz29) n=58. *mec-4*:HisCl +histamine n=20; no HisCl transgene +histamine n=11.

## Notes

### Competing Interest Statement

The authors have declared no competing interest.

### Summary of Updates

This version of the manuscript has been textually revised

